# Two-dye-imager DNA-PAINT enables volumetric nanoscopy of expanded cells

**DOI:** 10.64898/2026.05.18.725916

**Authors:** Janna Eilts, Julia Weingart, Cerridwen Kiesel, Harsha Perozhy, Philip Kollmannsberger, Arindam Ghosh, Dominic A. Helmerich, Markus Sauer

## Abstract

Refined single-molecule localization microscopy methods demonstrated superior localization precisions on isolated sample but remain limited by labeling density and imaging speed in cells. Here we combine expansion microscopy (ExM) with two-dye-imager (TDI)-DNA-PAINT to resolve fine molecular details of protein assemblies in ∼8-fold expanded cells with nanometer resolution. Using lattice light-sheet (LLS) microscopy, Ex-TDI-DNA-PAINT provides a robust platform for three-dimensional (3D) volumetric nanoscopy of the molecular organization of cells.

---

Single-molecule localization microscopy (SMLM) has transformed fluorescence imaging by enabling nanometer-scale spatial resolution imaging in cells^1^. Recent technical advances have substantially improved localization precision, in some cases approaching the Ångström regime for isolated fluorophores^2-4^. However, in cellular environments, achievable structural resolution is often limited by labeling density, probe accessibility, linkage error and imaging throughput rather than localization precision alone. Expansion microscopy (ExM), which physically enlarges biological specimens embedded in swellable hydrogels, has emerged as an alternative route towards nanoscale fluorescence imaging by increasing effective fluorophore distances and reducing the relative contribution of labeling offsets^5,6^. Post-expansion labeling has been proposed to improve epitope accessibility and reduce effective linkage error under suitable conditions^7-10^. Combining ExM with SMLM therefore represents a promising strategy for nanoscale imaging of molecular organization in cells^11-14^.

Early studies combining ExM with SMLM demonstrated the promise of this approach but also highlighted technical challenges associated with imaging expanded hydrogels^8,9,13,15,16^. These include reduced expansion homogeneity when preserving epitopes for post-expansion labeling, stabilization of highly expanded samples and compatibility with imaging buffers used for SMLM modalities.^15,16^ At the same time, current studies demonstrated that DNA Points Accumulation for Imaging in Nanoscale Topography (DNA-PAINT) based approaches are compatible with hydrogel environments^17^. Recently, a double-homogenized expansion microscopy workflow based on Ten-fold Robust Expansion (TREx) hydrogel chemistry was introduced for imaging of expanded cells at high spatial resolution^18^. In parallel, DNA-PAINT has become an important SMLM modality relying on transient interactions between fluorescently labeled “imager strands” and complementary “docking strands” attached to target molecules^19,20^. Since blinking is generated by programmable DNA binding kinetics, DNA-PAINT enables a complementary imaging strategy with flexible multiplexing capabilities^3,20^.

However, volumetric DNA-PAINT imaging of cellular samples is often limited by acquisition speed and background fluorescence arising from freely diffusing imagers^20-22^. To address these limitations, fluorogenic FRET-based and two-dye-imager (TDI)-DNA-PAINT imager strands were recently introduced^23-25^. Using TDI-DNA-PAINT, imager strands are labeled with two identical fluorophores that form quenched H-dimers in the unbound state, resulting in strongly reduced background fluorescence^23,24^. The fluorogenic properties of TDI strands permit imaging at increased imager concentrations and enable substantially faster acquisition compared with conventional DNA-PAINT^24^. Since expanded hydrogels represent optically large imaging volumes, we reasoned that fluorogenic TDI-DNA-PAINT could provide a suitable strategy for imaging expanded cellular samples.

To evaluate this possibility, cells were expanded following the previously described dTREx workflow (Supplementary Fig. S1)^18^. Briefly, cells were fixed using glutaraldehyde or formaldehyde, anchored into a TREx hydrogel^26^ and mildly denatured to facilitate post-expansion immunolabeling in partially expanded samples^18^. Following labeling with primary antibodies, samples were re-embedded into a second TREx hydrogel and subjected to proteinase K digestion to achieve approximately 8–9-fold expansion^18^. After immunostaining with secondary antibodies labeled with speed-optimized 7xR4 or 7xR3 docking strands^3,27^ at a degree of labeling (DOL) of 1-2, we re-embedded the sample into a neutral hydrogel to improve dimensional stability during imaging in high-salt DNA-PAINT buffers^12-14,18^.

In the first set of experiments, we immunolabeled microtubules with primary anti-α-tubulin antibodies and performed 2D Ex-TDI-DNA-PAINT imaging with R4 TDI strands labeled terminally with ATTO520 or ATTO Oxa14 dyes, respectively (Figure 1a)^24^. Microtubules are hollow cylindrical polymers assembled from α/β-tubulin heterodimers and represent established benchmark structures for super-resolution microscopy^28,29^. Super-resolved 2D images could be reconstructed already after 5-10 min acquisition time demonstrating the fast acquisition capabilities of TDI-DNA-PAINT (Supplementary Fig. S2). Expansion factors determined from microtubule diameter analysis using LineProfiler^12,30^ across multiple fields of view and cells, yielding an average expansion factor of ∼8.3 (Supplementary Fig. S3). The distribution was consistent across imaging sessions after re-embedding. Using highly inclined and laminated optical sheet (HiLO) illumination, we achieved theoretical and experimental localization precisions of 4.3 ± 2.6 nm (s.d.) and 5.65 ± 0.01 nm (s.d.) for ATTO520 and 5.1 ± 2.7 (s.d.) and 4.76 ± 0.01 nm (s.d.) for ATTO Oxa14, respectively^31,32^ (Supplementary Fig. S4). Three-dimensional biplane TDI-DNA-PAINT imaging of 8.3-fold expanded cells resolved the tubular organization of microtubules and enabled visualization of microtubule-associated labeling patterns in expanded samples^28,29,33^ (Figure 1b and Supplementary Fig. S5).

**Figure 1.**
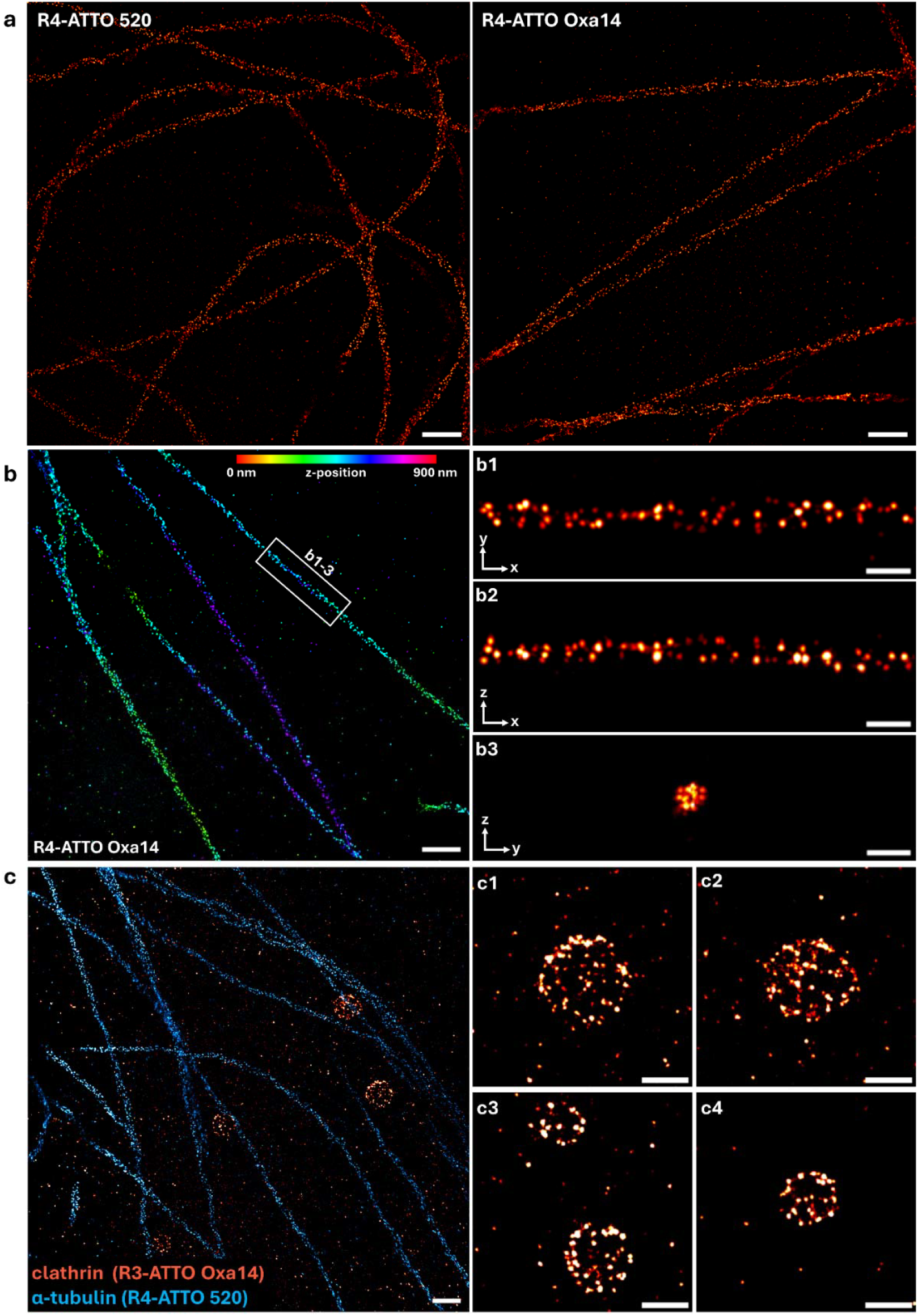
Ex-TDI-DNA-PAINT images of microtubules and clathrin coated pits in COS-7 cells. **a**, α-Tubulin was labeled with secondary antibodies carrying 7×R4 docking strands^3^. Measurements were performed in expanded samples using3.5 nM TDI strands R4-ATTO520 and R4-ATTO Oxa14, respectively, for 20 min. **b**, 3D-Biplane reconstruction of α-tubulin (TDI R4-ATTO Oxa14, 3.5□nM, 20□min). The color scale indicates z-position in expanded coordinates (0–900□nm). **b1**,**b2**,**b3**, Zoom-ins of white rectangle in (b) showing xy-, xz-, and yz-projections in expanded dimensions. **c**, Two-color Ex-TDI-DNA-PAINT image of α-tubulin (TDI R4-ATTO520, cyan) and clathrin heavy chain (TDI R3-ATTO Oxa14, red) acquired simultaneously at 3.5 nM imager concentration for 20□min. **c1-c4**, Enlarged views of representative clathrin-coated pits in expanded coordinates. Scale bars, 2□µm (a–c), 500□nm (b1–b3) and 1□µm (c1). All distances are shown in expanded sample coordinates unless otherwise stated.

Next, we performed multiplexing Ex-TDI-DNA-PAINT imaging of microtubules and clathrin-coated pits (CCPs) as representative cellular benchmarks. CCPs exhibit a diameter of 50-200 nm with characteristic polyhedral organization and are responsible for receptor-mediated endocytosis^34,35^. Therefore, we selected a second orthogonal docking strand (7xR3) which exhibits similar binding properties as 7xR4^27^. Using simultaneous imaging of TDI ATTO520-R4 and TDI ATTO Oxa14-R3 we were able to reconstruct multicolor Ex-TDI-DNA-PAINT images after acquisition times of only 10-15 min (Figure 1c and Extended Data Figure 1). The images enabled visualization of nanoscale features consistent with reported clathrin-coated pit morphology^35,36^.

Since TDI-DNA-PAINT imager strands are efficiently quenched in the unbound state we reasoned that they are ideally suited for volumetric 3D imaging using a commercially available inverted lattice light-sheet (LLS) microscope^22,24^. Single-molecule-sensitive volumetric imaging of 8.3x expanded cells was performed using high-power laser excitation and a light-sheet thickness of approximately 1.8 µm^24^. Three-dimensional localization was implemented using astigmatism introduced by laterally shifted aspheric plates^24,37^. Volumetric imaging of expanded cells spanning approximately 150 × 55 × 50 µm was completed within 13 hours (Figure 2a and Extended Data Figure 2). Reconstructed images show the 3D molecular organization of microtubules in cells with localization precisions of ∼20 nm in lateral and ∼45 nm in axial direction, which corresponds to effective localization precisions (divided by the expansion factor of ∼8.3) of ∼2.7 ± 1.8 nm (mean ± s.d.) in x-direction, 2.5 ± 1.6 nm in y-direction, and 4.5 ± 1.9 nm in z-direction (Figure 2b). In addition, physical expansion proportionally reduced the effective contribution of antibody linkage error and localization-cloud broadening associated with repetitive docking-strand designs^38,39^.

**Figure 2.**
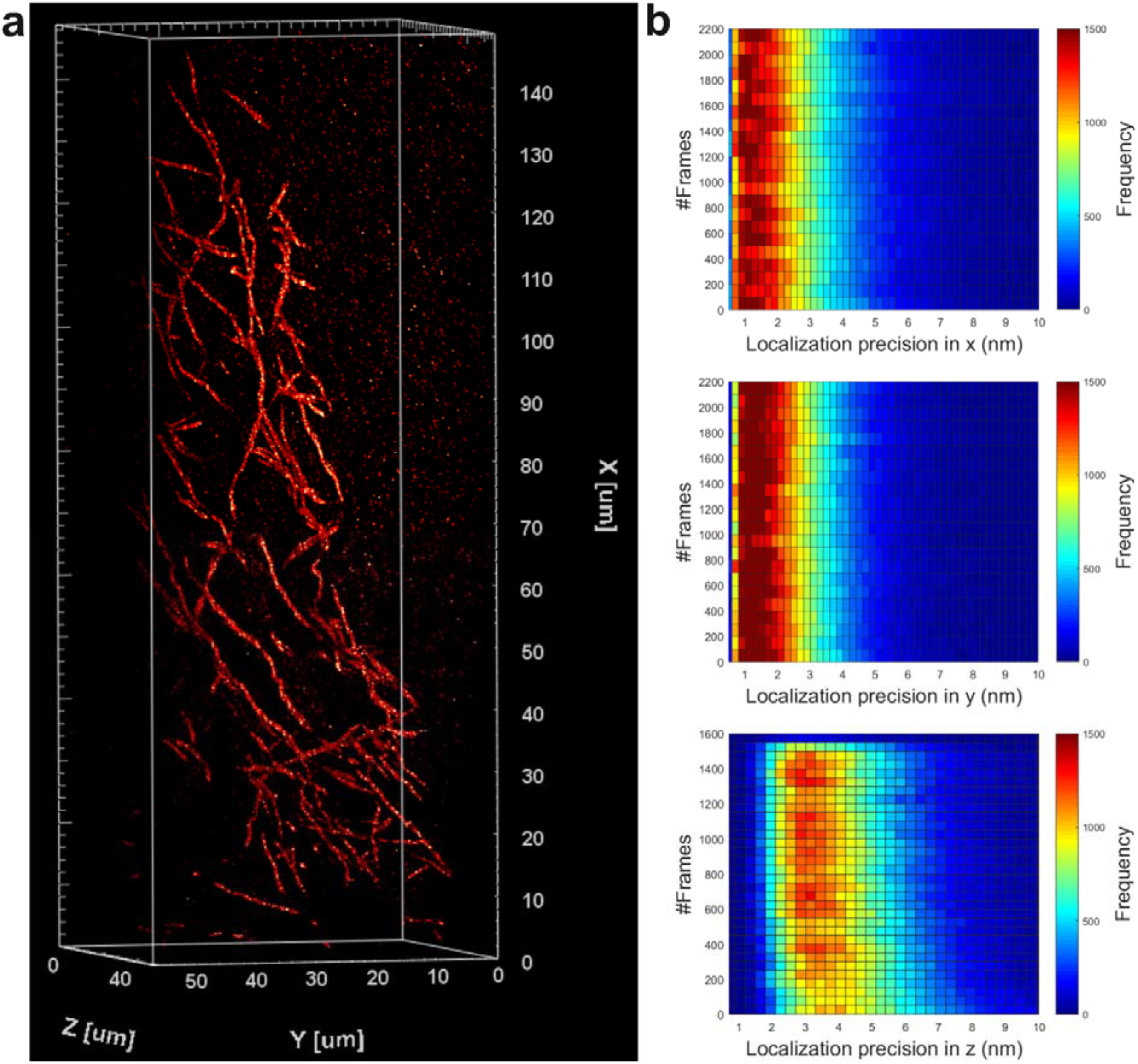
LLS-Ex-TDI-DNA-PAINT imaging of cellular microtubule networks. **a**, Whole-cell rendered volume 148 × 56 × 52 μm^3^ visualizing microtubule networks from an ∼8.3-fold expanded COS7 cell. **b**, Heatmap plots demonstrate effective localization precision values corrected for the expansion factor obtained from a representative stack, top panel, 2.7 ± 1.8 nm (mean ± s.d.) in x, middle panel, 2.5 ± 1.6 nm along y, and bottom panel, 4.5 ± 1.9 nm along z. Corrected effective localization precision values were calculated by dividing expanded localization precision by the mean expansion factor determined from microtubule diameter analysis.

Together, these experiments demonstrate that fluorogenic TDI-DNA-PAINT can be integrated with highly expanded cellular samples and volumetric lattice light-sheet imaging workflows as a complementary strategy. Our results establish Ex-TDI-DNA-PAINT as a proof-of-principle approach for combining fluorogenic DNA-PAINT with expansion microscopy for volumetric nanoscale imaging of cellular organization. Future integration with alternative fixation and preservation approaches, including cryofixation-based ExM workflows, may further extend the applicability of this strategy^40^.

## Methods

### Cell culture

COS-7 cells were cultured at 37°C with 5 % CO2 in DMEM/F12 medium (Merck #D8062) supplemented with 10 % FBS (fetal bovine serum, Merck #F7524) and 1% penicillin-streptomycin (Merck #P4333).

### Secondary antibody modification with docking strands

For labeling of goat anti-rabbit IgG (H+L) (Invitrogen #31212) and goat anti-mouse IgG (H+L) (Invitrogen #A28174) secondary antibody with PEG4-TCO, an excess of TCO-PEG4-NHS (Lumiprobe #3056) was used. Antibody labeling was performed at 20 °C for 4 h in labeling buffer (100 mM sodium tetraborate (Fulka #71999), pH 9.5). The antibodies were reconstituted in labeling buffer using 0.5 mL spin-desalting columns (40K MWCO, ThermoFisher #87766) following the manufacturer’s standard protocol. The TCO conjugated antibodies were purified using spin-desalting columns and stored in 1x PBS (PBS; Merck #D8537). The degree of labeling (DOL) was estimated by diluting TCO-modified antibodies in PBS, reacting them with an 8-fold excess of MeTet-dye via click reaction for 20 min at room temperature. Afterwards the samples were purified using 0.5 mL spin-desalting columns (40K MWCO, ThermoFisher #87766) to remove free dye and subsequently determining the dye-to-antibody ratio via UV/Vis absorption spectrometry. For TDI-DNA-PAINT measurements, modified goat anti-rabbit and goat anti-mouse antibodies were further modified with docking strands (7×R4 5’-3’: MetTet-ACACACACACACACACACA or 7xR3; 5’-3’: CTCTCTCTCTCTCTCTCTC) via click reaction for 20 min at 20 °C in PBS. The clicked antibodies were purified twice using 0.5 mL spin-desalting columns (40K MWCO, ThermoFisher #87766) and stored in PBS. Finally, the antibody concentration was determined by absorption measurements.

### *d*TREx with denaturation and digestion

Expansion gels were prepared according to the previously described *double*TREx (*d*TREx) protocol^17^. COS-7 cells were first fixed for 60 seconds at 37°C with 0.3 % glutaraldehyde (GA; Merck #G5882) and 0.25 % Triton™ X-100 (Triton; Merck #28314) in cytoskeleton buffer (CB-buffer). CB-buffer contained 10□mM 2-(N-morpholino)ethanesulfonic acid (MES; Merck #M3671), 150□mM NaCl (Merck #S7653), 5□mM ethylene glycol-bis(2-aminoethylether)-N,N,N′,N′-tetraacetic acid (EGTA; Merck #03777), 5□mM glucose (Merck #G7528) and 5□mM MgCl_2_; Merck #442615), adjusted to pH 6.1. Directly afterwards a second fixation step with 2 % GA in CB was conducted for 10 min. Cells were washed three times and optionally kept at 4°C. The anchoring step was done immediately before gelation with 0.25 % GA for 15 min. Samples were washed twice with PBS and once with TREx monomer solution^16^, consisting of 1.1 M sodium acrylate (Merck #408220), 2 M acrylamide (Merck #A4058), 0.009 % N,N’-methylenbisacrylamide (Merck #M1533), 1x PBS, 0.15 % Tetramethylethylenediamine (TEMED; Merck #T7024) and 0.15 % ammonium persulfate (APS; Merck #A7460). Coverslips (12 mm, round) were flipped on a drop of 60 µl TREx monomer solution with the cells facing the solution. Gels polymerized on ice for 15 min, then at room temperature for 1.5 h. Gels were homogenized in pre-heated denaturation buffer containing 200 mM sodium dodecyl sulfate (SDS; ThermoFisher #AM2548), 200 mM NaCl, 50 mM Tris (Merck, T1503) and 65 mM dithiothreitol (DTT; Merck #646563) (pH 8) for 1 h at 98°C. Five washing steps (15 min each) with warm PBS on a rotating wheel were performed before adding primary antibodies. Gel pieces (∼3 × 5 mm) were incubated with antibody solution overnight at 4°C and again with freshly prepared solution for 3 h at 37°C. For visualizing microtubules two primary antibodies were used in a mix (rabbit anti α-tubulin, abcam #ab18251 and mouse anti α-tubulin, Merck #T6199), each at a concentration of 10 µg/ml in 5 % bovine serum albumin (BSA; Merck #A7030) in each incubation step. For two-color measurements clathrin-coated pits were labeled with 15 µg/ml rabbit anti clathrin-heavy-chain (abcam #21679) and microtubules with 15 µg/ml mouse anti α-tubulin, Merck #T6199) in each step. After primary antibody incubation samples were washed with 0.1 % Tween^™^20 (ThermoFisher #28320) in PBS (PBST) on a rotating wheel for 3 × 15 min. A second anchoring step with 0.25 % GA for 20 min was then performed to anchor the primary antibodies into a second TREx gel. After three washing steps with PBS, each gel piece was incubated in 500 µl TREx monomer solution containing 0.03 % APS and 0.03 % TEMED two times for 30 min on a rotating wheel. During this step monomer solution diffused into the gels and caused them to shrink slightly to a ∼2.5x expanded state. Afterwards the monomer solution was removed and gels were “sandwiched” between to coverslips and polymerized in a humidified N_2_-filled chamber at 37°C for 1.5 h. Re-embedded gels were then homogenized a second time with a proteinase K digestion step. Therefore gels were incubated in digestion buffer consisting of 50 mM Tris, 1 mM EDTA (ethylenediaminetetraacetic acid disodium salt dihydrate; Merck #E1644), 0.8 M guanidine HCl (Merck #50933), 0.5 % Triton and 8 U/ml proteinase K (ThermoFisher #AM2548) for 2 h at 37°C. Samples were washed 5 x for 15 min with cold 0.1 % PBST and incubated with docking strand coupled secondary antibodies at a concentration of 20 µg/ml in 5 % BSA. Gels were incubated with antibody solution for 3 h at 37°C and additionally with freshly prepared antibody solution overnight. After three washing steps with 0.1 % PBST gels were fully expanded in a large volume ddH_2_O, before re-sembedding for TDI-DNA-PAINT imaging.

### Silanization of coverslips

For re-embedding and subsequent TDI-DNA-PAINT imaging, gels were mounted on silanized 24 mm round coverslips (1.5H, Carl Roth, PK26.2). Prior to silanization, coverslips were treated with three consecutive 15-minute washing steps in an ultrasonic bath: first with ddH_2_O, followed by 1 M KOH (Merck #P1767) and finally with 99 % ethanol. After drying, coverslips were incubated with silanization solution containing 0.001 % APTES ((3-aminopropyl)triethoxysilane; Merck #440140), 80 % ethanol (Merck #32205) and 0.02 % acetic acid (Merck #A6283) in ddH_2_O. Approximately 250 µl solution was applied to each coverslip and allowed to evaporate completely under a fume hood. Coverslips were then rinsed twice with 99 % ethanol, air-dried and stored at -20°C until further use.

### Re-embedding in neutral gel for TDI-DNA-PAINT imaging

To avoid shrinkage of expanded gels in ion-containing imaging buffer, samples were re-embedded in an uncharged polyacrylamide gel. All incubation steps were performed in 2 ml tubes placed on a rotating wheel. Initially, expanded gels were sectioned into thin slices (∼ 1 mm) with a razor blade to facilitate buffer diffusion. The slices were subjected to a crosslinking step with 0.25 % GA in ddH_2_O. for 20 min, followed by three washes of 15 min each in ddH_2_O. Subsequently, gels were incubated twice for 45 min in re-embedding solution consisting of 10 % acrylamide, 0.15 % Bis, 0.025 % TEMED and 0.025 % APS in ddH_2_O. After removing excess solution, the gels were transferred onto silanized 24 mm coverslips with the cell side facing the coverslip. A second untreated coverslip was placed on top. Polymerization was carried out for 1.5 h at 37-40°C in a humidified box filled with nitrogen.

### DNA-PAINT image acquisition

Re-embedded gels were incubated in imaging buffer (1x PBS + 500 mM NaCl, pH 7.4) for 30 min and immersed in fresh imaging buffer containing imager strands at a concentration of 3.5 nM 15 min before imaging. Measurements were performed on an inverted fluorescence widefield microscope (Olympus, IX-71) equipped with a nosepiece stage (Olympus, IX2-NPS) and two EMCCD cameras (Andor iXon Ultra DU-897). The localization precision was calculated according to Mortensen et al.^30^ and Endesfelder et al.^31^. Biplane 3D images were acquired with an 60x oil objective (Olympus, NA 1.45 PlanApo) and HILO (highly inclined and laminated optical sheet) illumination using ∼ 1 kW/cm^2^ of an appropriate laser (Coherent, Genesis MX 639). For recording on two EMCCD cameras (Andor iXon Ultra DU-897) simultaneously, a two-channel image splitter (TwinCam, Cairn Research) equipped with an 50/50 beamsplitter (Cairn Research) was used and the cameras were synchronized by a pulse generator (DG535, Stanford Research Systems) Subsequently, frames were acquired on both cameras with a frame rate of 10 Hz for 15-30 min. A dichroic mirror (ZT405/514/635rpc, Chroma) and a bandpass filter (Em01-R442/514/647-25, Semrock) was placed in front of the objective to separate excitation and emission light. In addition, a quarter-wave plate (Thorlabs, SAQWP05M) was mounted for excitation with circular polarized light. For calibration, 100 nm microspheres (Invitrogen, T7279) were prepared by incubating ∼105 beads/ml in 1xPBS with 50 mM MgCl_2_ adjusted to pH 7.4, for 15 min using eight chambered cover glass systems with high performance cover glass (Cellvis, C8-1.5H-N), followed by three washing steps prior calibration experiments. The calibration measurements were performed by using a piezo scanner (Pifoc, Physik Instrumente) driven with a LVPZT servo controller (E-662, Physik Instrumente) to move the objective. For two color DNA-PAINT imaging an oil-immersion objective (APON 60×, numerical aperture 1.49; Olympus) was employed. Samples were excited by a 639 nm laser and 514 nm laser (Genesis MX 639-1000 STM and Genesis MX 514-500 STM, Coherent) respectively. A dichroic mirror (ZT405/514/635rpc, Chroma) and a bandpass filter (Em01-R442/514/647-25, Semrock) was used to separate excitation from emission light. A beamsplitter (630 DCXR customized, Chroma) projected emission light through two different bandpass filters (582/75 and 679/41 BrightLine series, Semrock) on two separate EMCCD cameras. The two channels were acquired simultaneously in HILO illumination with an exposure time of 100 ms/frame (10 Hz) and an irradiation intensity of ≤ 1 kW/cm^2^ of the 514 nm laser and the 639 nm laser. Images were reconstructed in the open-source software rapidSTORM 3.3^39^ with a pixel size of 20 nm. For two-color alignment 0.2 µm fluorescent beads were imaged in each channel and an alignment matrix was created with the ImageJ plugin bUnwarpJ^40^. The matrix was later applied to the reconstructed images.

### Three-dimensional biplane analysis

For extracting 3D information, an intensity-based analysis routine was used. First, the measured image stacks were analyzed with rapidSTORM 3.3^39^. Therefore, the FWHM was set to 360□nm, the intensity threshold was set to 250 photons and the fit window radius to 1100□nm. The corresponding localization files from both cameras were then analyzed further using a custom written python script to calculate the 3D intensity ratios. The calibration file as well as the sample files measured, were analysed in the same way as described elsewhere in detail but focused on the rapidSTORM analysed intensity values for calculating the intensity ratios^41,42^. For visualizing 3D images, ImageJ (version 2.16.0/1.54g) and ThunderSTORM (version 1.3) was used.

### Lattice light-sheet microscopy

For PSF-calibration 100 nm fluorescent beads (Invitrogen #T7279) were diluted 1:400 in 1 % agarose (Sigma-Aldrich #A9539) and the agarose was allowed to polymerize on a glass slide. A piece of the bead-containing agarose gel was subsequently positioned on the silanized coverslip adjacent to the re-embedded sample gel. The sample gel was incubated in imaging buffer (1x PBS + 500 mM NaCl, pH 7.4) for 30 min and immersed in fresh imaging buffer containing imager strand R4-Oxa14 at a concentration of 0.5-1 nM 15 min before imaging. LLS 3D TDI-DNA-PAINT experiments were performed as described by Ghosh et al.^22^. In brief, imaging was carried out on a commercial lattice light-sheet microscope (Lattice Lightsheet 7, Zeiss) equipped with a 13.3×/0.4 NA excitation objective and a 44.83×/1.0 NA detection objective, and an ORCA-Fusion sCMOS camera (Hamamatsu), yielding an effective size of 145 nm in the specimen plane. Only the 640 nm excitation line was employed at a nominal power density of 3.3 kW/cm^2^ at the light sheet. Volumes were recorded with 150 ms exposure time and typically 2,500 frames per slice. To enable 3D single-molecule localization, astigmatism was introduced via the built-in aberration control. Fiducial beads were acquired in a separate volumetric scan (0.2 µm stage steps in the skewed y-direction) and later used to generate an averaged 3D PSF and an astigmatism-based z-calibration curve.

Microscope control, volumetric acquisition, on-the-fly deskewing and automated stepping along the skewed y-axis were handled by the vendor software (ZEN) using a custom macro adapted^24^ that defined the stage step size (typically 0.8 µm), number of slices and file naming. During acquisition, the macro logged both nominal and actual stage positions to a.csv file, which was subsequently used for precise geometric deskewing and alignment of slices.

Single-molecule analysis followed the lattice light-sheet TDI-DNA-PAINT workflow established elsewere. Deskewed calibration stacks were processed in FIJI (ImageJ) using DoM for bead detection and PSF extraction, followed by iterative PSF averaging with RegisterNDFFT to obtain a high-signal 3D PSF and a corresponding z-calibration that accounts for astigmatism and lateral wobble. Experimental time series were then batch-processed with custom FIJI macros calling DoM to perform single-molecule localization (typically assuming a PSF standard deviation of 1.8 – 2.0 pixels and a signal-to-noise threshold of 4–8), assign z-coordinates via the stored calibration, correct intra-series drift and generate per-slice super-resolution reconstructions together with localization tables. Per-slice reconstructions were deskewed using the recorded stage positions and registered against one another by cross-correlation to correct residual inter-slice drift, following the strategy outlined in Ghosh et al.^24^. Finally, all localization tables were merged in MATLAB/Python scripts adapted from the original analysis code to apply the measured geometry, drift corrections and rotation (typically 30°) into a coverslip-aligned Cartesian coordinate system. The resulting combined localization table contained 3D coordinates (X, Y, Z in nm) and localization uncertainties and was used for volumetric rendering and quantitative analysis. All macros and scripts were directly derived from or minimally adapted versions of the publicly available TDI-DNA-PAINT analysis pipeline accompanying Ghosh et al.^24^.

## Estimation of expansion factor

For estimating the expansion factor, the average diameter of expanded microtubules was determined multiple fields of views using the LineProfiler software^25^. Therefore, images of isolated single strands were cropped out of overview 2D Ex-TDI-DNA-PAINT images to avoid incorrect placements of line profiles over two crossing or closely spaced strands. Configuration parameters in the software for the proper detection of linear structures were set in the following ranges: Gaussian blur 70-100; spline parameter 2.0-3.0. However, these parameters do not influence the original image and measured distances. In summary 24 ROIs in 8 overview images of 3 independent experiments were analyzed, after discarding ROIs where many lines were not set correctly. The 24 line profiles (resulting from in total 8053 set lines) were normalized, averaged and fit with a bigaussian function, yielding a peak-to-peak distance of ∼207 nm. Divided by the known diameter of unexpanded microtubules of 25 nm this results in an expansion factor of ∼8.3. A small number of incorrectly set lines and the linkage error is neglected as the antibodies could potentially bind to the inside and outside of the microtubules in post-labeling.

## Extended Data Figures

**Extended Data Figure 1.**
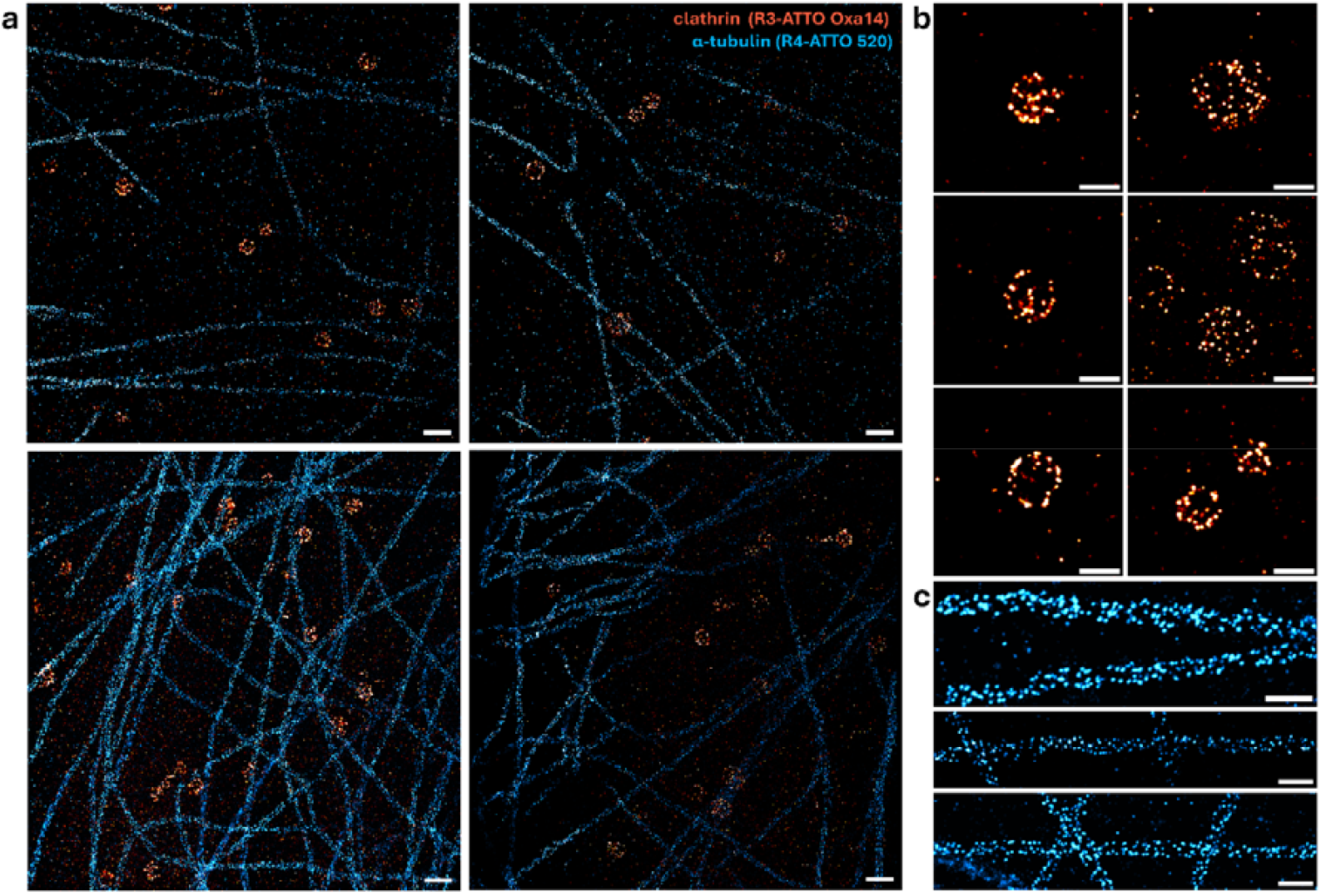
Two-color Ex-TDI-DNA-PAINT of microtubules and clathrin coated pits (CCPs) in COS-7 cells. **a**, Two-color images of α-tubulin visualized by TDI strand R4-ATTO 520 (cyan) and clathrin heavy chain by TDI strand R3-ATTO Oxa14 (red) in expanded samples. **b**,**c** Zoomed-in images of CCPs (b) and microtubules (c) in expanded coordinates. Scale bars, 2 µm (a), 1 µm (b,c) correspond to expanded sample dimensions.

**Extended Data Figure 2.**
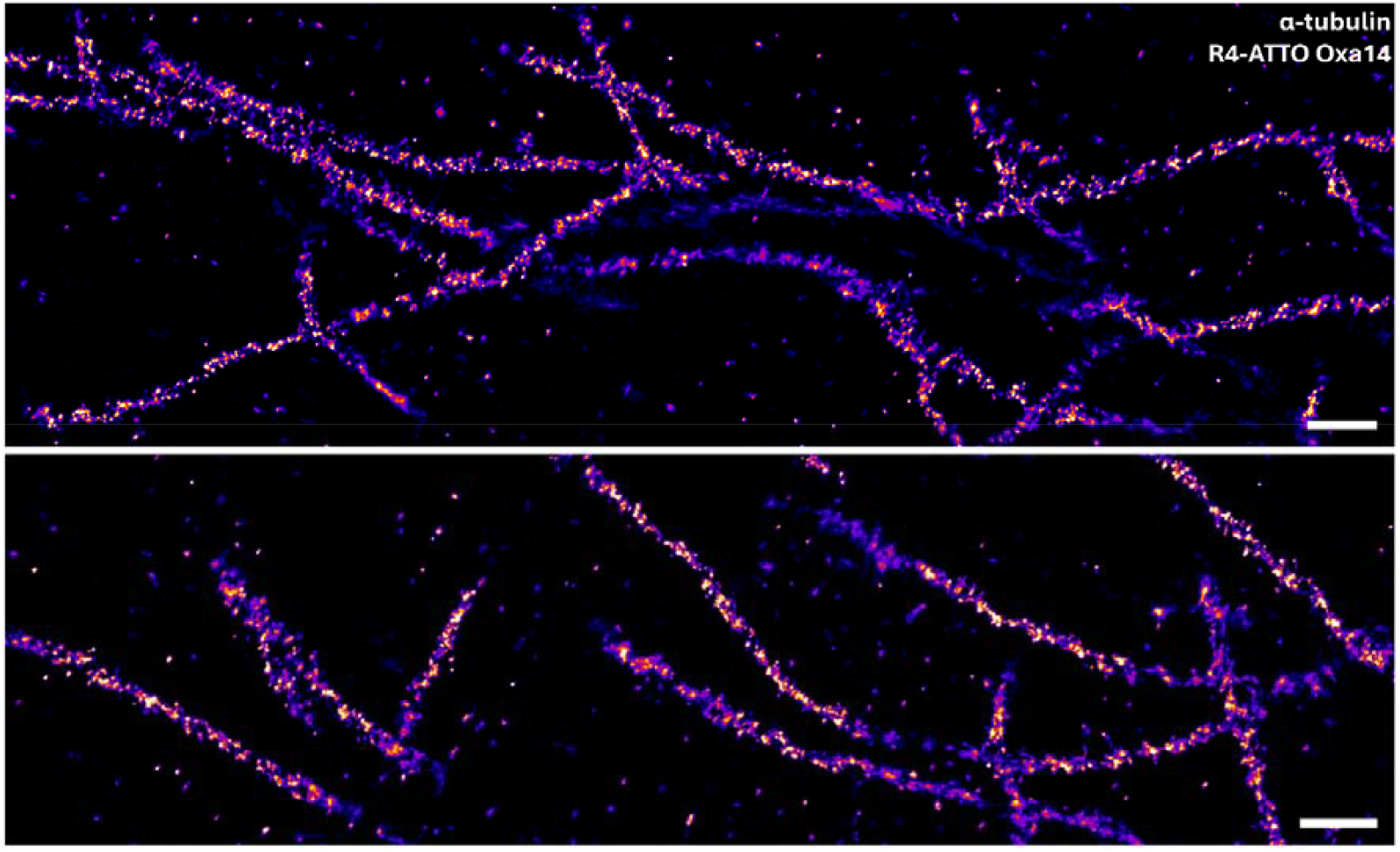
Lattice light-sheet Ex-TDI-DNA-PAINT of microtubules in COS-7 cells. Representative 2D slices from deskewed 3D-stacks of α-tubulin imaged in ∼8.3× expanded samples using TDI R4-ATTO Oxa14. All images are shown in expanded coordinates unless otherwise stated. Scale bars, 2 µm.

