## Supplementary Figures for "Two-dye-imager DNA-PAINT enables volumetric nanoscopy of expanded cells"

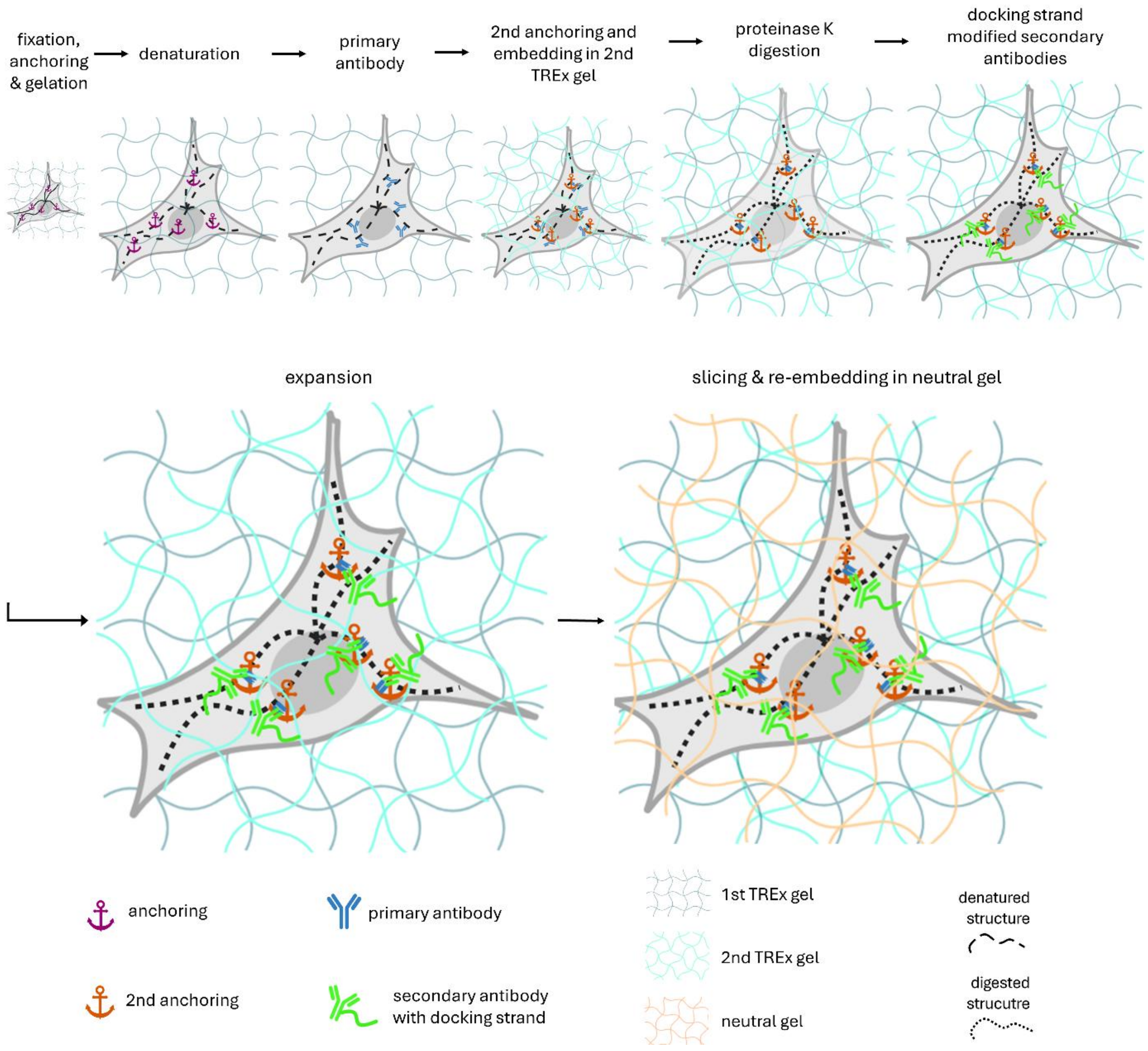


**Supplementary Figure 1. Sample preparation for TDI-DNA-PAINT imaging using *double*TREx (*d*TREx)[1].** Fixed cells are anchored into a TREx[2] gel. After initial mild homogenization using denaturation with heat and detergent, primary antibodies are applied. Partly expanded gels (~2.5-fold) are then embedded into a second TREx gel, in which the primary antibodies are now anchored. This enables a subsequent harsh proteinase K digestion to completely homogenize the sample. Fragments of primary antibodies which are crosslinked in the second gel can then be addressed by secondary antibodies conjugated with docking strands before gels are fully expanded in deionized water. Prior to imaging in salt-containing TDI-DNA-PAINT imaging buffer, gels are sliced into ~ 1 mm thin sections and re-embedded into an uncharged polyacrylamide gel to prevent gel shrinkage during imaging. Created with Biorender.com.

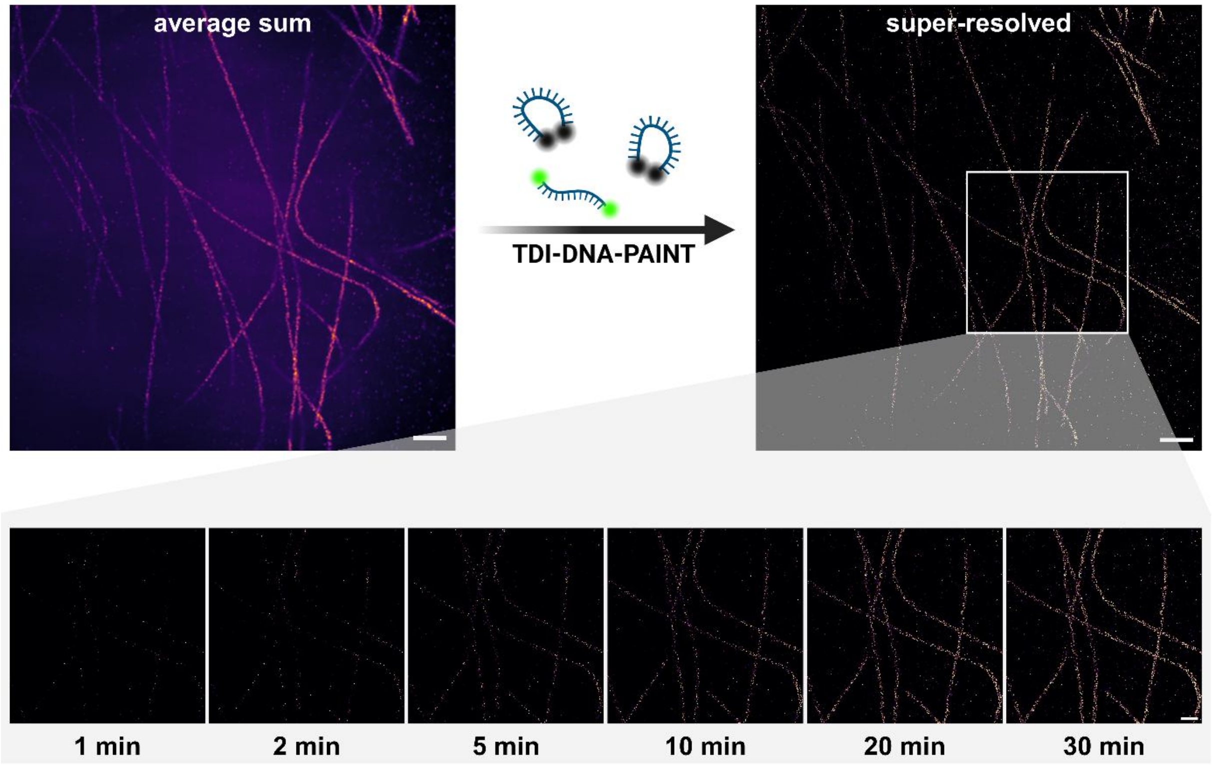


**Supplementary Figure 2. Ex-TDI-DNA-PAINT reconstructions of microtubules in COS-7 cells after different acquisition times.** Averaged fluorescence image of all recorded frames (top left) and corresponding TDI-DNA-PAINT super-resolution reconstructions (top right). The zoomed-in region below shows reconstructions obtained after 1, 2, 5, 10, 20, and 30 min of recording. High-quality reconstructions with clearly resolved filament structures are already achieved after 10–20 min, demonstrating the fast acquisition capability of TDI-DNA-PAINT. Scale bars, 5 µm (top) and 2 µm (insets).

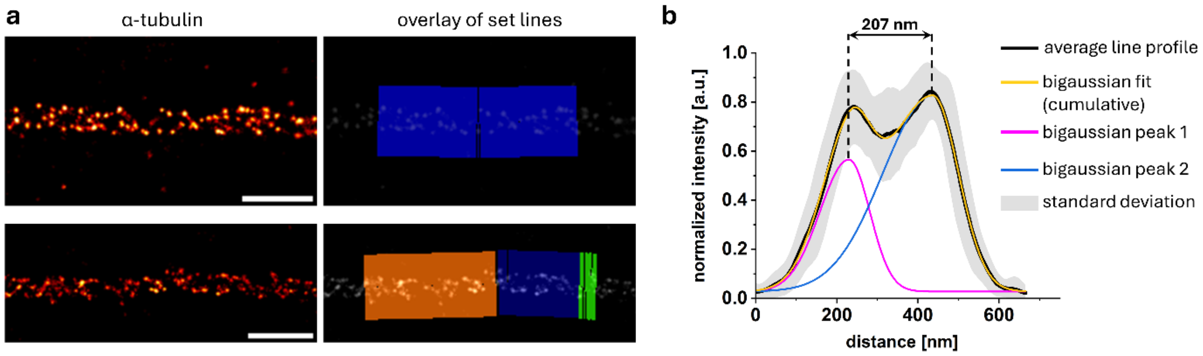


**Supplementary Figure 3. LineProfiler[3] analysis for estimation of the expansion factor. a**, Exemplary regions of interest (ROIs) used for analysis with the LineProfiler software, showing isolated filaments from Ex-TDI-DNA-PAINT images of α-tubulin (left). Overlay of lines set by the LineProfiler shows accurate detection of the filament (right). **b**, Averaged line profile (black) from 24 ROIs. Fitting with a bigaussian function yielded a cumulative peak fit (yellow) and separate peak fits for peak 1 and peak 2. The distance between peak fit 1 and peak fit 2 was used to determine the average diameter of the expanded microtubules of 207 nm. Divided by the known diameter of unexpanded microtubules of 25 nm, this results in an expansion factor of ~8.3-fold. Scale bars, 1 μm (expanded dimensions).

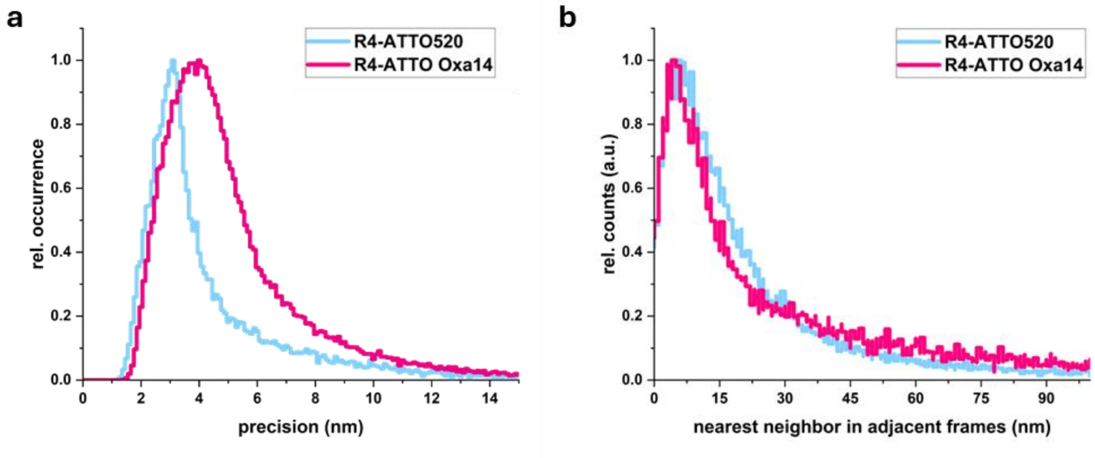


**Supplementary Figure 4. Localization precision analysis of R4 TDI-DNA-PAINT imagers.** **a**, Localization precision of ATTO520 (cyan) or ATTO Oxa14 (magenta) modified R4 imagers based on photon counts per localization[4]. **b**, Experimental localization precision of the same R4 imagers determined by nearest-neighbor analysis[5]. Scales are not corrected for the expansion factor.

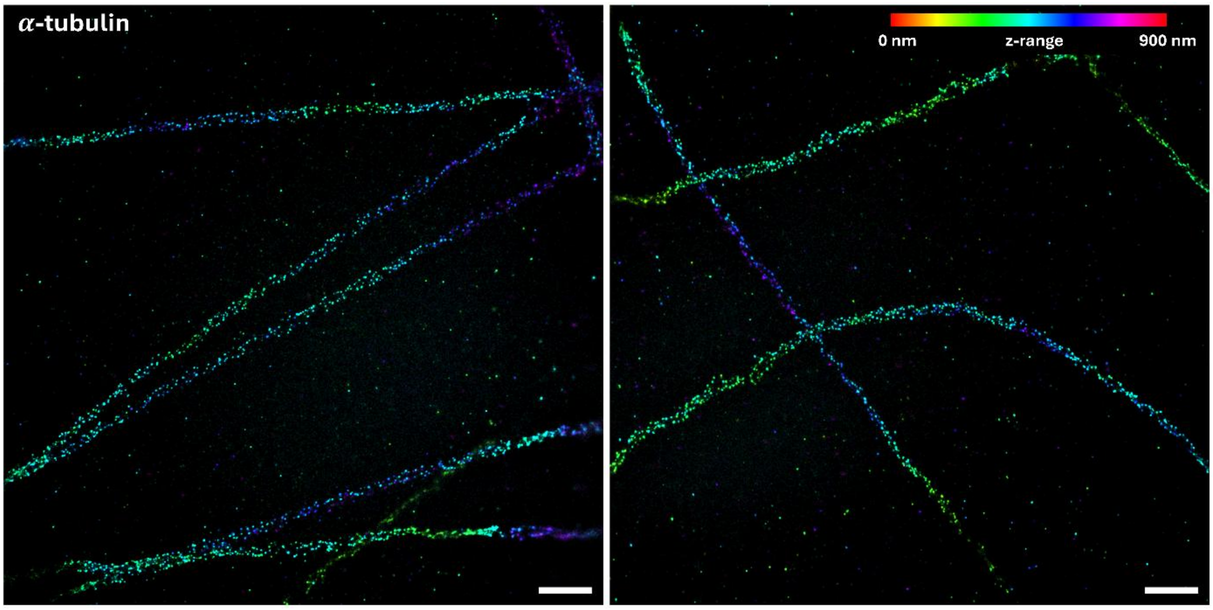


**Supplementary Figure 5. 3D Ex-TDI-DNA-PAINT of microtubules.** Additional color-coded 3D biplane reconstructions of microtubules (α-tubulin) in COS-7 cells obtained using Ex-TDI-DNA-PAINT. The color scale represents the axial position of localizations from 0–900 nm. The images reveal highly resolved filament structures extending across different depths, showcasing the capability of Ex-TDI-DNA-PAINT for volumetric nanoscale imaging. Scale bars, 2 µm (expanded dimensions).

**Description of Supplementary Videos**

**Supplementary Video 1.** Ex-TDI-DNA-PAINT recording of immunostained microtubules using imager strand R4-Atto 520 (corresponding to reconstructed two-color image in Figure 1c). Frames 1-2000 are shown at a rate of 10 frames per second (fps).

**Supplementary Video 2.** Ex-TDI-DNA-PAINT recording of immunostained clathrin heavy chain using imager strand R3-Atto Oxa14 (corresponding to reconstructed two-color image in Fig. 1c). Frames 1-2000 are shown at a rate of 10 fps.

**Supplementary Video 3.** Single plane of 3D Ex-TDI-DNA-PAINT recording using lattice light-sheet (reconstructed 3D projection in Figure 2). Immunostained microtubules are visualized with imager strand R4-Atto Oxa14. Frames 1-2000 are shown at a rate of 10 fps.

**Supplementary Video 4.** Reconstructed 3D Ex-TDI-DNA-PAINT lattice light-sheet stack (corresponding to 3D projection in Figure 2). Immunolabeled microtubules are visualized with imager strand R4-AttoOxa14.
